# Binary Relation Extraction from Biomedical Literature using Dependency Trees and SVMs

**DOI:** 10.1101/082479

**Authors:** Anuj Sharma, Vassilis Virvilis, Tina Lekka, Christos Andronis

## Abstract

The goal of Biomedical relation extraction is to uncover high-quality relations from life science literature with diverse applications in the fields of Biology and Medicine. In the last decade, several methods can be found in published literature ranging from binary to complex relation extraction. In this work, we present a binary relation extraction system that relies on sentence level dependency features. We use a novel approach to map dependency tree based rules to feature vectors that can be used to train a classifier. We build a SVM classifier using these feature vectors and our experimental results show that it outperforms simple co-occurrence and rule-based systems. Through our experiments, using two ‘real-world’ examples, we quantify the positive impact of improved relation extraction on Literature Based Discovery.

## 1 INTRODUCTION

In silico drug repositioning [1] seeks new indications for existing drugs based on Computational Biology and Computational Chemistry methods. Conceptually, in silico drug repositioning focuses on quantifying the space between drugs and disease or its molecular targets. Databases and corpora such as PubMed have been used extensively for drug repositioning via text mining [2]. Well studied methods based on Information Retrieval and Information Extraction [3] have been used for mapping documents to a lower dimensional concept space for semantic analysis. Based on this type of analysis, non-obvious or even unknown (hence, novel) relations between a drug and a disease may then be uncovered.

Techniques for identification of biomedical relations broadly belong to two categories: interaction based methods and text mining methods [4]. The former use data from “biomedical interaction” databases such as such as Genes-to-Genes (e.g. STRING [5]), Genes-to-Chemical compounds (e.g. STITCH [6]), Genes-to-Diseases (e.g. OMIM [7]) and other more general repositories (e.g. IntAct [8]). The types of biological relations in text can be a) semantic, b) grammatical, c) negation and coreference [15]. This work focuses on identifying semantic relations among biological entities.

In addition to the interaction databases, many (if not more) biomedical interactions are also described, albeit in the form of free-text, in scientific publications. PubMed/Medline [30] is the largest and most comprehensive biomedical database containing more than 21 million (as of late 2014) biomedical abstracts. Medline is increasing by over 700,000 abstracts every year. The availability of biological knowledge in the form of free text (biomedical literature), has motivated many research groups to apply text mining methods in order to “extract” biomedical relations from free text [11]. Text mining approaches vary from simple co-occurrence [12] - where two entities are considered to be related if they occur in the same abstract, to those using sophisticated Natural Language processing (NLP) [13] and machine learning [14] techniques. Excellent reviews of relation extraction methods can be found in [11], [15], [16].

The NLP-based approaches differ from one another in several aspects. One of them is the technique employed to analyze the input text, i.e. Part-of-speech (POS) tagging [17], shallow parsing [18] or full parsing [19] but also in the methods for extracting and learning rules [11].

In general, a relation extraction system using NLP consists of: a) Named Entity Recognition (NER) [28], b) Relation Trigger Word Identification and c) Relation Extraction [31]. Each of these tasks is non-trivial presenting challenges such as polysemy, affecting both NER and identification of trigger words, owing to which several approaches have been proposed for each of them [15]. While relation trigger word identification is typically used in relation extraction it is not a mandatory step and maybe left out in generalized relation extraction systems. A pre-processing step may also be included in order to facilitate the remaining steps. Preprocessing module segments the document into basic text units and augments the text for subsequent processing.

Several studies in the biomedical domain target binary relation extraction due to its potential to enhance our knowledge of protein-protein interactions [20], [21], [22], [27], [29]. Most such systems assume normalized entity names and employ dictionary based approaches for the NER step [11]. The methods belong to two categories: a) rule-based approaches and b) machine learning-based approaches. Typically textual analysis such as POS tagging, syntactic parsing and dependency parsing is applied in the sentences. A combination of features is then extracted and represented as a rule or used for training a classifier. Mostly, supervised machine learning methods have been employed although some attempts have been made to use semi-supervised and unsupervised machine learning techniques [4]. Support Vector Machines (SVMs) have been used with great success in machine learning based binary relation extraction techniques [11], [23].

In this work, we present a sentence level binary relation extraction system based on SVMs. Biological entities were identified using a dictionary and pairs of entities were classified by a SVM as related or unrelated to each other (per sentence). We developed a novel approach for representing a pair of entities, occurring in a sentence, as a feature vector constructed from dependency tree based rules. We used these feature vectors for building a classifier and compared its performance to that of simple rule-based and co-occurrence based methods. We also quantified the impact of improved relation extraction in a “real world” scenario using our Literature-Based Discovery platform [31], [32], [33], [34], [35].

## 2 METHODS

### 2.1 Binary relation extraction

The binary relation extraction process developed in this work is shown in Fig. 1. A trained classifier is built using a corpus where sentences and related entities appearing in them are known. A sentence containing a pair of proteins can then be classified, as implying an interaction or a lack thereof using this trained classifier. It should be noted that a sentence containing *n* entities will be processed 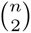 times (combinatorial) i.e. once for each pair. The sentence must first be pre-processed to ensure that the entities can be correctly identified (*Named Entity Recognition*). This is followed by generating a dependency parse tree for the sentence. Finally, feature vectors are extracted from the tree for classification.

**Fig. 1.**
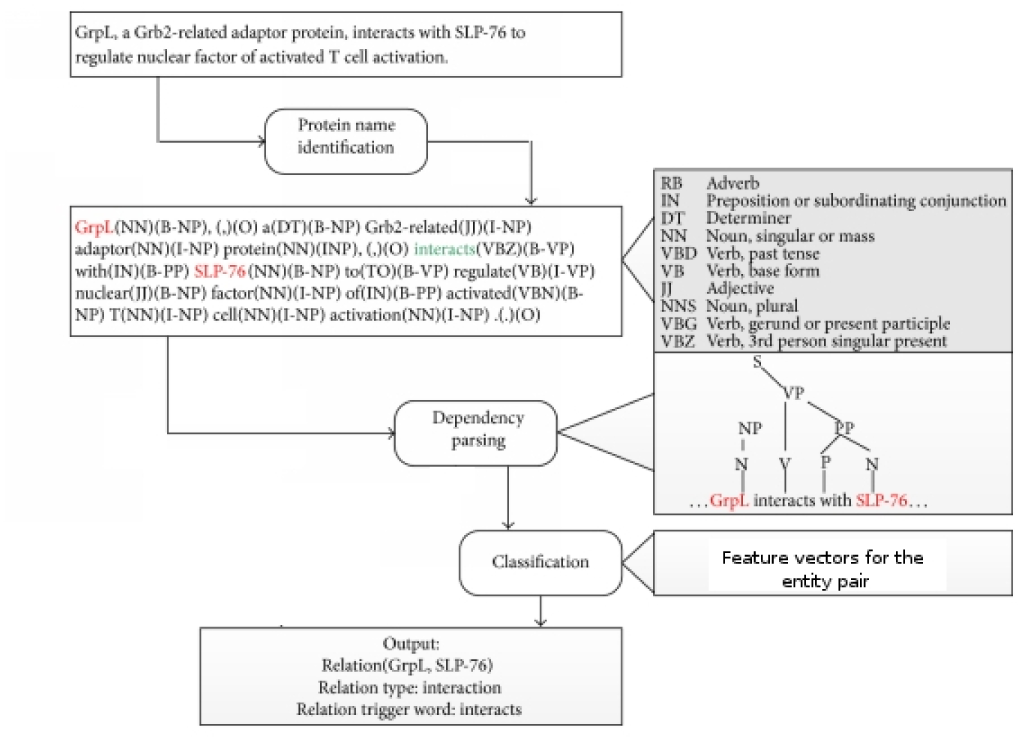
**Overview of binary relation extraction process developed in this work. A sentence is first pre-processed to ensure the entities contained can be identified, this is followed by generation of a dependency parse tree for the sentence. Subsequently paths relating pair of entities extracted and feature vector constructed for classification, adapted from [11].**

#### Named entity recognition

To extract relations between entities, it is important to correctly identify the entities. However, this is not a trivial task due to issues such as polysemy and non-standard gene/protein nomenclature used in the scientific text [11]. We use a dictionary-based approach to address the problem. A list of entity names expected to appear in the corpus, ensures that all entities are found during parsing. Further, to avoid segmentation of multi-part entity names during dependency parsing we pre-process the corpus replacing space in such names with underscores (giving preference to longer names). This is a basic heuristic that facilitates the automated extraction task without leaving out the entities that are described by lengthy clusters of noun phrases. This straightforward approach to pre-processing safeguarded that the entries of interest have been correctly recognized and tagged. There was only one reservation about this annotation option, namely that the ‘shrinking’ of sentence parts (especially NP phrases) might be counter-effective

#### Automatic rule extraction

We refer to a sequence of tags along the branches of a dependency tree as a rule. Figure 2 shows a dependency tree for a sentence. Using the tree the rule that can be extracted showing a relation between *NF-kappa_B* and *I_kappa_B_beta_inhibiter is: prep_of:amod::prep_by*, *by*, with *Control* being the starting point. Given a sentence and entities contained it is possible to determine the rules (paths) that associate these entities using the dependency tree for the sentence. We developed a tool to automatically extract such rules for all pairs of entities in a given sentence (from its dependency tree). Repeating this process over the entire corpus results in a set consisting of rules that a) apply to entity pairs that are related and b) entity pairs that are unrelated. The final set consists of all unique rules of the former type and all unique rules of the latter type. It is clear that the set will contain some overlapping rules (those that associate both related and unrelated entities). We use this set as the universe of complex linguistic rules (Ξ). The rules extracted can themselves be used for rule-based identification of relations between entities in a give sentence.

**Fig. 2.**
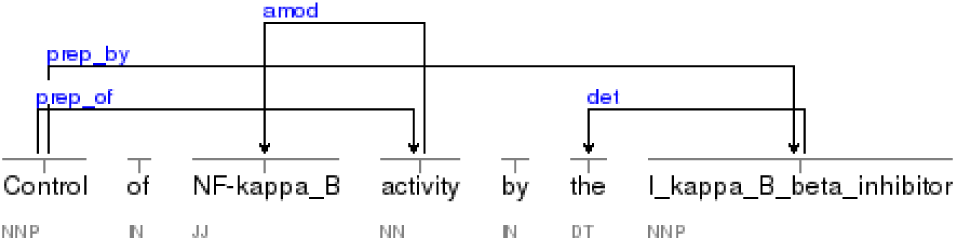
**The dependency tree generated for a sentence can be used for associating biological concepts of interest. In this sentence its clear that there exists a path between *NF-kappa_B* and *I_kappa_B_beta_inhibiter*.**

#### Feature extraction and building a classifier

Using Ξ each pair of entities can be represented by a feature vector of *N* dimensions, where *N* = cardinality of Ξ. Conversion of an entity pair to feature vector is carried out as follows. A dependency tree is generated for the sentence where the pair of entities appears. The tree is used to extract all rules associating the pair of entities. A binary vector of dimension *N* can then be generated by placing 1 for all rules in Ξ that are found for this pair of entities. Thus a sparse vector representation, based on dependency parsing, is generated for each entity pair for each sentence. A training set generated using this procedure, consisting of feature vectors for related entities and unrelated entities, can be used to train a classifier.

### 2.2 Application to Literature based discovery (LBD)

With a view of carrying out a ‘real-world’ evaluation of the relation extraction process we integrate it to Biovista’s LBD platform and compare the performance of the vanilla and enhanced versions [35]. The LBD platform allows a user to uncover indirect associations between biological concepts such as Diseases, Drugs and Adverse Events giving the user options on how the associations are scored and what associations are considered at each stage. Further, the output list is sorted by the strength of the associations, reflecting the relevance of each concept to those input biological concepts. We integrated the binary relation extraction process to the LBD platform, allowing replacement of the co-occurrence based list with a list generated by the relation extraction process.

### 2.3 Datasets

The Genia Meta-Corpus dataset [26] was used as the basis for creating the GMCR dataset used in this work to benchmark the relation extraction process. The corpus consists of 1000 MEDLINE abstracts with manually annotated biomedical events and has been extensively used for benchmarking Information Extraction systems. The Genia dataset contains annotations per sentence which define the entries in the text and correlate them with events. Detailed description of the format can be found online at http://www.nactem.ac.uk/genia/genia-corpus. Briefly, the entries (word or group of words) in a sentence are separated into terms (Ts) and non-terms (As). The terms typically refer to biological concepts while non-terms may or may not belong to this category. Events, belonging to a short list of events selected as relevant to biomedicine, define causes and theme relationship.

Figure 3, shows an entity-relationship graph generated from a sentence taken from the Genia Meta corpus. A cause (theme) can be a term or another event. Using this information we extracted pairs of terms (biological concepts) that are related to one another. For example sentence 1 of abstract with pubmed id 10022882 - *“Reactive oxygen intermediate-dependent NF-kappaB activation by interleukin-1beta requires 5-lipoxygenase or NADPH oxidase activity”* - consisting of annotations shown in Table 1. From the figure relations extracted would be T1-T2, T3-T2 and T4-T2. Table 2 shows the size of the resulting dataset.

**Fig. 3.**
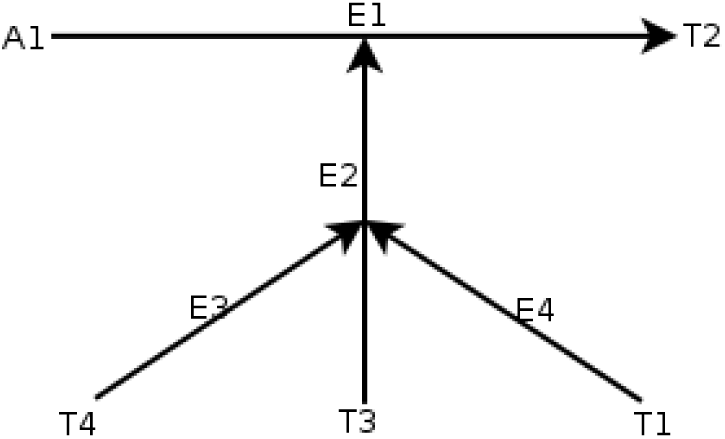
**Sample event tree generated from Sentence 1 of annotated abstract with Pubmed Id 10022882 in the Genia Meta Corpus.**

**TABLE 1.** Tags and associated terms extracted from Genia Meta Corpus for the first sentence of the abstract with pubmed if 10022882.

| Tag | Term in text |
| --- | --- |
| A1 | Reactive oxygen intermediate |
| T1 | NADPH oxidase |
| T2 | NF-kappaB |
| T3 | Interleukin-1beta |
| T4 | 5-lipoxygenase |

**TABLE 2.** **Size of the pre-processed dataset generated from GENIA meta-corpus to maximize NER.** The pre-processing applied to the corpus replaces spaces in all multi-part entity names with underscores. This ensures that when the corpus is parsed the entities are identified correctly. Nested entities and associated relations are lost because the pre-processing is biased towards longer entity names.

|  |  |
| --- | --- |
| Total number of abstracts | 1000 |
| Total number of unique sentences | 9372 |
| Total number of relations | 6735 |

Two datasets, listed in Table 3, were used for quantifying the impact of improving relation extraction on literature based discovery. Both datasets comprise commonly used drugs for Alcohol addiction and Multiple Sclerosis. Drugs are presented by their active ingredients. Drug names were extracted from free-text using all their synonyms, including brand names to increase recall at the NER step.

**TABLE 3.**
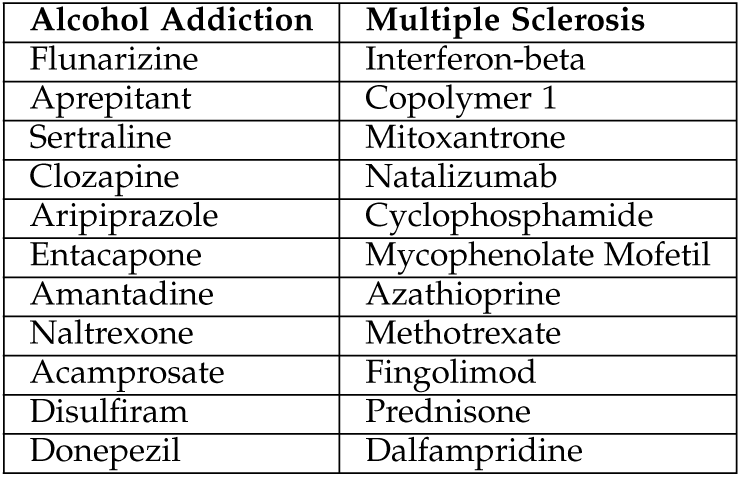
**Dataset with Alcohol addiction used for quantitative impact of relation extraction on literature based discovery.**

| Alcohol Addiction | Multiple Sclerosis |
| --- | --- |
| Flunarizine | Interferon-beta |
| Aprepitant | Copolymer 1 |
| Sertraline | Mitoxantrone |
| Clozapine | Natalizumab |
| Aripiprazole | Cyclophosphamide |
| Entacapone | Mycophenolate Mofetil |
| Amantadine | Azathioprine |
| Naltrexone | Methotrexate |
| Acamprosate | Fingolimod |
| Disulfiram | Prednisone |
| Donepezil | Dalfampridine |

## 3 RESULTS AND DISCUSSION

### 3.1 Experimental system used

All experimental results presented in this work were generated on a Intel Xeon X5660 CPU running at 2.80GHz with 24Gb of RAM. The processor contains 6 cores hyper-threaded to 12 and each machine has two such processors. Due to the large volume of data processing required (specifically for the LBD step) two such machines were used for distributing various tasks.

### 3.2 Automatic extraction of rules and Feature vector generation

We extracted linguistic rules from the GMCR dataset relating biological concepts from sentence dependency trees generated using Stanford-Core NLP [24]. Based on the principles of unsupervised grammar induction, the Stanford Dependency Parser (SDP) operates on the notion of dependency, i.e. on the idea that the syntactic structure of a sentence consists of binary asymmetrical relations between the words of the sentence. In addition, it is a parsing system that emphasizes the importance of local syntactic context, thus making it useful for describing the interaction between entities. This set captures rules which result in valid as well as invalid associations between the concepts. This process resulted in generation of a rule set consisting of 89528 rules. A binary feature vector, for a pair of entities in a sentence, of dimension 89528 can then be generated by placing 1 at indexes corresponding to rules that appear in the dependency tree of the sentence. A classifier trained with features based on such a rule set must therefore also be able to find the combination of such rules that correctly identifies relations.

### 3.3 Performance comparison of SVM classifier with cooccurrence

In order to benchmark the binary relation extraction process we compared it with a co-occurrence based co-relation extraction system. The simple co-occurrence based system assumes that when two entities appear together in an abstract they are necessarily related to each other. A natural consequence of this simple assumption is that, when named-entity recognition (NER) is assumed to have Precision and Recall equal to 1, the co-relation extraction system has Recall equal to 1.0 since all possible relations are captured. However, a side effect of this is high False Positive count lowering the Precision of the overall system. Nevertheless, such a system serves as an excellent baseline system, because a subject matter expert (e.g., a Biologist or Clinician) using the information extracted is likely to prefer higher False Positives than False Negatives. Further, we also compared the performance of using the rules extracted with the classifier.In order to use the rules without a classifier a pair of biological concepts with contain a path in the dependency tree (of the sentence) that appears in the rule set are assumed to be related.

Figure 4, shows the results of SVM parameter analysis. We used a subset of the dataset, with equal number of positive and negative feature vectors (10% of the positive feature vectors), to find the best combination of SVM parameters. In order for the classifier to be generalizable it is ideal that it has balanced performance on both the classes (positive and negative) in terms of accuracy. The figure shows that a balanced performance across four metrics - Precision, Recall, F-score and True Negative Rate (Accuracy on −1) - was achieved in our experiments at a value of g = 0.01, c = 100 and w = 2. We also carried out 10 iterations of evaluation with the most balanced classifier, with the test set containing being unique from the training set, to not the average performance of the SVM classifier on the four metrics. Table 4, shows the average performance of the classifier with the mean and standard deviation of the values obtained. As can be seen the classifier performance does not have a large variation which is a desirable characteristic. Subsequently, a leave-one-out analysis of the classifier (over the entire dataset) yielded Precision of 0.99 and Recall of 0.16. The performance was attributed to the large imbalance between the negative and positive feature vectors in the training set (15:1). If a more balanced dataset is used the performace expected would be closer to that obtained in Table 4.

**Fig. 4.**
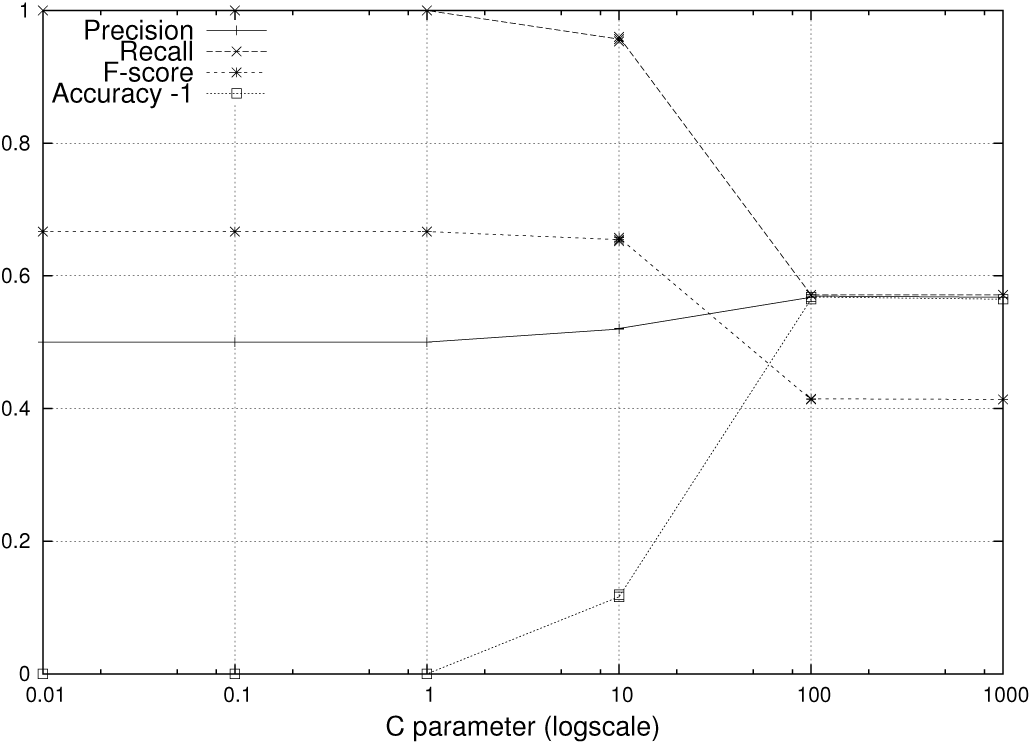
**Variation of F-measure, Precision (P), Recall (R) and Accuracy on −1 with change in c parameter of the SVM (g = 0.01 and w = 2). The best performance for a balanced classifier is achieved at c = 100.**

**TABLE 4.** **Mean and standard deviation of SVM performance, g = 0.01 c = 100 and w = 2, in terms of Precision, Recall, Accuracy (Acc.) on +1 (identifying sentence with a relation) and Accuracy on −1 (identifying sentence without a relation) calculated over 10 iterations**

|  | Precision | Recall | Acc. (+1) | Acc. (-1) |
| --- | --- | --- | --- | --- |
| Mean | 0.59 | 0.49 | 0.49 | 0.64 |
| Standard Deviation | 0.08 | 0.11 | 0.11 | 0.14 |

Table 5 shows a comparison of the SVM classifier with those of the co-occurrence and simple rule-based systems. As expected for the co-occurrence based system the Recall is very high because any pair occurring together in a sentence is considered to be related, however this comes at the cost of the Precision of such a system. The rule-based relation identification, based on the Auto-rule extraction, performs slightly better than simple co-occurrence in terms of Precision but has very low Recall. This is because such a relation identification system has no generalization and only features that have been seen in the training set can be correctly identified in the test set. However, the SVM classifier is capable of learning and generalizing and out-performs both the simplistic baseline systems. Finally, while it is difficult to compare different relation-extraction schemes, due to the differences in specifics and datasets, our method copmares favorably with those of other state-of-the-art methods with an F-score of around 0.50 as presented in [11].

**TABLE 5.** **Comparison of relation extraction performance between simple co-occurrence, rule based extractor using the automatically extracted rules and the SVM.** Co-occurrence based relation extraction correctly identifies all relations but has the drawback of having very low precision. The Auto-rule based relation extraction improves the precision while retaining the recall. The SVM clearly has the best performance on the comparison metrics.

| Method | Precision | Recall | TNR | F-measure |
| --- | --- | --- | --- | --- |
| SVM | 0.59 | 0.59 | 0.60 | 0.59 |
| Auto-rules | 0.04 | 0.1 | 0.99 | 0.06 |
| Co-occurrence | 0.01 | 1.0 | - | 0.02 |

### 3.4 Qualitative analysis of classifier based relation-extraction

Finally we compare the performance of the NLP advanced LBD system with the base system on a real world dataset. The starting point for the experiment is the Alcohol Addiction disease (Multiple sclerosis disease) and the required output is a ranked list of drugs likely to be related to the disease. Table 6 and Table 7 show the performance comparison between the different LBD setups. It can be seen that as expected the SVM based LBD outperforms the basic LBD and the LBD with the auto rule set. It was interesting to note that the performance of the SVM based filtering scheme andthe auto-rules were comparable. This suggests the high-quality of the rule set that was extracted by the automatic procedure. Clearly the rule set provides sufficient diversity to capture previously unseen relations. However, the SVM based approach is likely to be more generalizable and therefore it is likely to be the better approach to use.

**TABLE 6.** **Relative ranks of drugs in LBD results with Alcohol Addiction as the starting entity.** Each column shows the rank of the drugs in LBD results when the first-level expansion is filtered by the specific relation extraction method (co-occurrence, auto-rule and SVM). Cells with the best rank achieved per row are highlighted.

| Drug | Base LBD | Auto-rule | SVM |
| --- | --- | --- | --- |
| flunarizine | 1429 | 502 | 1274 |
| aprepitant | 1392 | 1448 | 1208 |
| sertraline | 286 | 129 | 96 |
| clozapine | 66 | 100 | 116 |
| aripiprazole | 184 | 135 | 84 |
| entacapone | 1202 | 1973 | 1930 |
| amantadine | 179 | 211 | 303 |
| naltrexone | 26 | 6 | 2 |
| acamprosate | 394 | 14 | 8 |
| disulfiram | 34 | 2 | 6 |
| donepezil | 696 | 289 | 408 |

**TABLE 7.** **Relative ranks of drugs in LBD results with Multiple Sclerosis as the starting entity.** Each column shows the rank of the drugs in LBD results when the first-level expansion is filtered by the specific relation extraction method (co-occurrence, auto-rule and SVM). Cells with the best rank achieved per row are highlighted.

| Drug | Base LBD | Auto-rule | SVM |
| --- | --- | --- | --- |
| Interferon-beta | 2 | 1 | 1 |
| Copolymer 1 | 15 | 3 | 2 |
| mitoxantrone | 196 | 6 | 12 |
| natalizumab | 25 | 9 | 7 |
| Cyclophosphamide | 18 | 17 | 84 |
| Mycophenolate Mofetil | 369 | 69 | 52 |
| Azathioprine | 271 | 34 | 27 |
| Methotrexate | 109 | 33 | 30 |
| fingolimod | 24 | 5 | 4 |
| prednisone | 139 | 22 | 25 |
| dalfampridine | 3163 | 27 | 15 |

These experiments have shown that some of the true positives (e.g. good drug candidates for repositioning) may be found as “low” as position 1000. While this number might seem large, it is still the top 1000 of 20,000 possible drugs, in other words it represents the top 5% of the list. Furthermore since many of the better-placed candidates may be clear false positives, we feel that the burden on the subject matter expert of shifting through the top 5% of a list is usually well worth the effort. Put more precisely, this decision may ultimately be left to the user who will act taking into account the stakes at hand. If the “lower hanging fruit” are suitable and available from a discovery and IP perspective, then the experts effort may be limited to the top 2–3% of each list. If however the “lower hanging fruit” are not available, then the end user may choose to extend his effort to the top 5–6% of the ranked list with a good chance of finding useful correlations.

The above results suggest that false positives in correlations diminish the performance of the predictive platform and reduce the sensitivity towards the different parameter combinations. Effort should be therefore expended in reducing the number of false positive in correlations for example through the use of NLP techniques.

## 4 CONCLUSION

In this work we present a novel approach for sentence-based Binary relation extraction from Biomedical literature (specifically from PubMed abstracts). We have developed a novel method for mapping rules extracted from the dependency tree of the sentence to feature vectors that can be used for training and classification using an SVM classifier. Our results, show that the SVM classifier achieves an F-score of 0.59 outperforming simple rule-based (F-score of 0.06) and co-occurrence (F-score of 0.02) based systems. Further, we show that such a classifier can have a high impact on the performance of a Literature Based Discovery system in terms of associating diseases and drugs by presenting two “real world” examples.

Precise automated relation extraction systems in combination with high-throughput Literature Based Discovery (LBD) platforms can reduce the amount of data required to be manually analyzed by subject matter experts. In a discovery set-up where it is often more important to ensure that “no stone is left unturned” we believe that this man-machine relationship will serve the purpose better than one that may offer better precision but at the expense of missing discovery opportunities. Furthermore in the specific case of where the raw materials (i.e. our knowledge of biology and disease) are incomplete and often conflicting, building biological causation arguments that incorporate multiple causal paths may be a prudent approach to utilize.

Due to the positive outcome observed in our experiments, we plan to extend our work, through the use of more sophisticated NLP tools such as Anaphora, Nominalization and Negation to increase Recall. More experimental evaluation will be required to study the impact this would have on the overall classifier performance. Another direction of work that we intend to investigate is qualifying the relations by way of recording trigger words or other context words. This would allow deeper analysis into types of relations that are problematic and perhaps use ensemble of classifiers each dedicated to a different type of relation.

## ACKNOWLEDGMENTS

This work was supported by the European Union ICT Program (Project “p-medicine from data sharing and integration via VPH models to personalized medicine” FP7-ICT-2009.5.3, #270089).

The authors wish to acknowledge Drs. Spyros Deftereos, SVP, Drug Discovery at Biovista and Eftychia Lekka, Senior Investigator at Biovista for their help with the sets of Drugs chosen for Alcohol Addiction and Multiple Sclerosis and also their overall contribution as subject matter experts in Drug Repositioning.

## Anuj Sharma

Anuj Sharma is currently working as a Data Scientist in the innovation team for EXUS. He was a staff scientist and developer at Biovista with expertise in Ontologies and Literature mining research.

## Tina Lekka

Tina Lekka is an independent scholar, currently working on a book on Conceptual Metaphor Theory. Her PhD research was conducted at University College London (UCL) and her academic interests lie in the fields of Textual Analysis and Cognitive Linguistics.

## Vassilis Virvilis

Vassilis Virvilis is Head of IT at Biovista. He has a PhD in Artificial Intelligence. He has a long standing interest in Machine Learning Algorithms and Optimization.

## Christos Andronis

Christos Andronis is Vice President for Research & Development at Biovista. He has a Ph.D. in Biochemistry from Imperial College, London, UK. He has been involved in the design and implementation of Information Extraction systems and other Computational applications relevant to Drug Discovery.

